# RNAP II produces capped 18S and 25S ribosomal RNAs resistant to 5’-monophosphate dependent processive 5’→3’ exonuclease in polymerase switched *Saccharomyces cerevisiae*

**DOI:** 10.1101/2021.06.28.449823

**Authors:** Miguel A. Rocha, Bhavani S. Gowda, Jacob Fleischmann

**Affiliations:** Research Division, Greater Los Angeles VA Healthcare System, Los Angeles, California, USA; David Geffen School of Medicine at UCLA, Los Angeles, California, USA; School of Dentistry at UCLA, Los Angeles, California, USA

## Abstract

We have previously found in the pathogenic yeast *Candida albicans*, 18S and 25S ribosomal RNA components, containing more than one phosphate on their 5’-end, resistant to 5’-monophosphate requiring 5’→3” exonuclease. Several lines of evidence pointed to RNAP II as the enzyme producing them. We now show in *Saccharomyces cerevisiae*, permanently switched to RNAP II, due to deletion part of RNAP I upstream activator alone or in combination with deletion of one component of RNAP I itself, the production of such 18S and 25S rRNAs. They contain multiple phosphates at their 5’-end and an anti-cap specific antibody binds to them indicating capping of these molecules. These molecules are found in RNA isolated from nuclei, therefore are unlikely to be capped in the cytoplasm. This would be unlike recapping of decapped mRNAs which occurs in the cytoplasm. Our data confirm the existence of such molecules and firmly establish RNA II playing a role in their production. The fact that we see these molecules in wild type *Saccharomyces cerevisiae* indicates that they are not only a result of mutations but are part of the cells physiology. This adds another way RNAP II is involved in ribosome production in addition to their role in the production of ribosome associated proteins.

## Introduction

Eukaryotic cells devote a large percentage of their energy resources to the production of ribosomes (Warner, 1999), the protein producing organelles located in the cytoplasm. They are made up of structural and synthetically active RNAs combined with over 70 proteins (Woolford and Baserga, 2013). In yeast, the generation details of the rRNA components are well established. The genes coding for rRNAs are grouped in tandem repeats separated by non-transcribed spacer sequences (NTS) (Thiry and Lafontaine, 2005). The NTS contains the rDNA promoter with its upstream element (UE) and core element (CE) representing the initiation site of rDNA transcription (Goetze et al., 2010). This transcription requires the binding of upstream activating factor (UAF), a multiprotein complex consisting of Rrn5, Rrn9, Rrn10, Uaf30, histones H3 and H4, to the upstream element and TATA binding protein (TBP) (Keener et al., 1997); (Siddiqi et al., 2001). The transcription of rDNA is carried out by RNA polymerase I (RNAP I) resulting in a 35S rRNA precursor molecule processed into 18S, 25S and 5.8S rRNA components (Planta and Raue, 1988);(Haag and Pikaard, 2007). The gene for the fourth component 5S, is located within the NTS, and transcribed by RNA polymerase III (RNAP III) in the reverse direction (Turowski and Tollervey, 2016). These components are mostly assembled with the ribosomal proteins in the nucleus and are exported and completed in the cytoplasm (Kargas et al., 2019). The genes coding for the ribosomal proteins are transcribed by RNA polymerase II (RNAP II) thus giving all three RNA polymerases a role in ribosome biogenesis (Abraham et al., 2020).

Since ribosomal RNA (rRNA) represents over 80% of total RNA produced by cells, a major function for rRNA has become to serve as control for quality and quantity of RNA isolation, for studies that focus on the many coding and non-coding smaller RNAs (Schroeder et al., 2006). 5’-monophosphate dependent 5’ to 3’ processive exonucleases, such as Terminator (Lucigen), have become available, to enhance the recovery of RNAs of interest, by eliminating the dominating rRNA from total isolated RNA. The utility of these enzymes is based on the fact that processed RNA molecules typically have a single phosphate on their 5’-end, making them vulnerable to these enzymes (Jager et al., 2009). Capped RNAs typically are protected from digestion, aiding in their recovery. In studies utilizing Terminator involving the polymorphic yeast *Candida albicans*, we unexpectedly found that this yeast was producing 18S and 25S rRNA components resistant to digestion as it shifted to the stationary growth phase (Fleischmann and Rocha, 2018). Digestion of rRNA with tobacco acid pyrophosphatase made these 18S and 25S molecules again susceptible to Terminator digestion. This indicated that the 5’-ends of these Terminator resistant molecules, contained more than one phosphate. Additional studies with the same yeast, that included RNAP I inhibition, chromatin immune precipitation with anti-RNA II antibodies and immunoblotting with anti-cap specific antibodies were carried out (Fleischmann et al., 2020). The sum of these experiments indicated that in addition to its role in ribosomal protein production, RNAP II plays a role in the production of these exonuclease resistant RNA molecules.

While typically RNAP II is suppressed from gaining access to the rDNA promoter site, allowing RNAP I an exclusive role in rRNA transcription, in *Saccharomyces cerevisiae* a role for RNAP II in rRNA production is well established (Vu et al., 1999); (Vallabhaneni, 2015). This yeast contains tandem repeats of rDNAs of 9.1-kb length on chromosome XII as many as 200, but can also have a 9.1-kb monomer episomal circles of rDNA (Clark-Walker and Azad, 1980). They are excisional products of homologous recombination between tandem repeats. Such an episomal circular rDNA containing respiratory deficient *Saccharomyces cerevisiae* with a cryptic RNAP II promoter has been described (Conrad-Webb and Butow, 1995). This organism could utilize RNAP II to generate a 35S precursor off the episomal rDNA circle, while possibly utilizing RNAP I for copying off the tandem repeats. Similarly, mutants with *UAF30* deletion also utilize both RNAP I and RNAP II for rRNA transcription (Siddiqi et al., 2001). Deletion of *rrn9*, one of the UAF components results in complete switching to RNA II, designated as a PSW phenotype (Oakes et al., 1999). Additional deletion of RPA 135 component of RNAP I in these PSW yeast, confirmed the sole role of RNAP II in rRNA transcription in these cells (Yano and Nomura, 1991) (Nogi et al., 1991). Primer extension studies of the 5’-end of the polycistronic 35S precursor molecules showed them to be variable from-9 to −95 from the known RNAP I promoter site, suggesting a separate or overlapping promoter for RNAP II in these mutants. Recently, we became aware of the availability of such PSW phenotypes of *Saccharomyces cerevisiae* from YGRC/NBRP in Japan, which allowed us to test the validity of RNAP II’s role in the production of exonuclease resistant 18S and 25S rRNA components.

## Methods

### Organisms

*Saccharomyces Cerevisiae* S288C (ATCC), BY27539 (*MAT**a** ade2-1 ura3-1 his3-11 trp1-1 leu2-3,112 can1-100 rrn9*Δ*::HIS3 rpa135*Δ*::LEU*) and BY27384 (*MAT**a**/a ade2-1/ade2-1 ura3-1/ura3-1 his3-11/his-3-11 trp1-1/trp1-1 leu2-3,112/leu2-3,112 can1-100/can1-100 RRN9/rrn9*Δ*::HIS3*) (YGRC/NBRP Japan) were maintained in 50% glycerol in YPD broth (2% w/V tryptone, 1% w/v yeast extract, 2% w/v dextrose) at −80°C. Cells were activated in YPD broth at 30°C and maintained on Sabouraud dextrose agar at 4°C, passaged every 4–6 weeks up to 4–5 times. Yeasts were lifted from agar surface and grown in YPD broth for variable length of times at 30°C. Yeast cell concentrations were established using a hemocytometer.

### RNA Isolation

Cells were collected by centrifugation, washed with sterile phosphate buffered saline (PBS) and were put on ice pending total RNA extraction. Cells were disrupted with RNase-free zirconia beads and RNA was isolated using Ambion RiboPure RNA Purification kit for yeast (Ambion/ThermoFisher) according to the manufacturer’s instructions.

Nuclear RNA was obtained using the Yeast Nuclei Isolation kit (Abcam) following the manufacturer’s instructions. Histone Acetyltransferase (HAT) Activity Assay Kit (Abcam) was used to verify nuclear source of isolated RNA. RNA quantification and quality were assessed by using a Qubit 4 fluorometer and an Agilent 2100 Bioanalyzer.

### Terminator 5’-Phosphate-dependent Exonuclease Experiments and 5’-end analysis

Total RNA was treated with Terminator (Lucigen) following the manufacturer’s protocol using the supplied Buffer A. The ratio of enzyme to substrate employed was 1 U per 1 μg of RNA to ensure adequate cleavage. RNase inhibitors were used in all the assays.

### RNA Analysis

Terminator treated and non-treated RNA samples were loaded into an RNA 6000 Nano chip and analyzed with the Agilent 2100 bioanalyzer system (Agilent Technologies, INC). Electropherograms were used to calculate Terminator resistance percentages by measuring the areas under the peaks of untreated (uncut) RNA and dividing it by the area of treated (cut) RNA.

### Immunoblotting

RNA was separated on formaldehyde agarose gels (Lonza) and stained with SYBR Gold Nucleic Acid Gel Stain (Life Technologies) for 30 minutes. Gel images were captured with a digital camera (Canon Vixia HFS30). RNA was transferred by electro-blotting (BIO-RAD Trans-Blot Turbo Transfer System) to a positively charged nylon membrane (Millipore) in 0.5 x TBE (standard Tris/Borate/EDTA buffer). The RNA was cross-linked to the membrane using UV (Stratagene UV Crosslinker). Membrane was blocked with 10% Block Ace™ (Bio-Rad) for 30 minutes at 25°C, followed by the addition of anti-m3G-cap, m7G-cap antibody clone H20 (Millipore Sigma) diluted 1:1000 in 10% Block ACE™ and incubated for 24 hours at 4°C. Goat anti-mouse conjugated to HRP was added to the membrane at 1:5000 in blocking solution for 30 minutes at 25°C. The Prosignal ™ (Prometheus) chemiluminescence substrate was used to detect the HRP signal. Film was developed with the SRX-101A Konica film processor.

### Decapping Assays

Cap-Clip™ acid pyrophosphatase (Cellscript) was used according to manufacturer instructions for decapping RNA samples. Verification of cap removal was done by gel electrophoresis, Northern blotting and immunoblotting using anti-cap (H20) antibody. Biotinylated probes used for Northern blot were made using the following sequences: 18S_3_Fwd (5’-GTGAAACTCC GTCGTGCTGGG-3’), 18S_3_Rev (5’-TAATGATCCTTCCGCAGGTTCAC CTAC-3’), 25S_3_Fwd (5’-AACGCGGTGATTTCTTTGCTCCAC-3’), 25S_3_Rev (5’-GGCTTAATCT CAGCAGATCGTAACAACAAGG-3’)

## Results and Discussion

To buttress the evidence derivable from the single and double mutants we needed to establish the behavior of wild type *Saccharomyces cerevisiae* regarding 18S and 25S molecule resistance to Terminator digestion. As can be seen in Figure 1, Terminator eliminates 18S and 25S rRNAs completely from total RNA isolated from wild type yeast cells during active growth period. As the cells approach the stationary growth phase, 18S and 25S rRNAs resistant to Terminator begin to appear. This can be seen directly in stained gels (Fig. 1A), as well by Northern blotting (Fig. 1B). Assaying these molecules through a Bioanalyzer (Figure 1C-D) not only confirmed their presence also but allowed us to quantitate them (Fig. 1E). About 30% of the 18S and 25S RNAs become resistant to Terminator digestion during stationary growth phase. The fact that both total and nuclear RNA result in the same percentage, indicates that they are produced in the nucleus and not recapped in the cytoplasm. Thus, the wild type *Saccharomyces cerevisiae* mimics the pattern of Terminator resistance we observed in *Candida albicans* (Fleischmann and Rocha, 2018).

**FIGURE 1.**
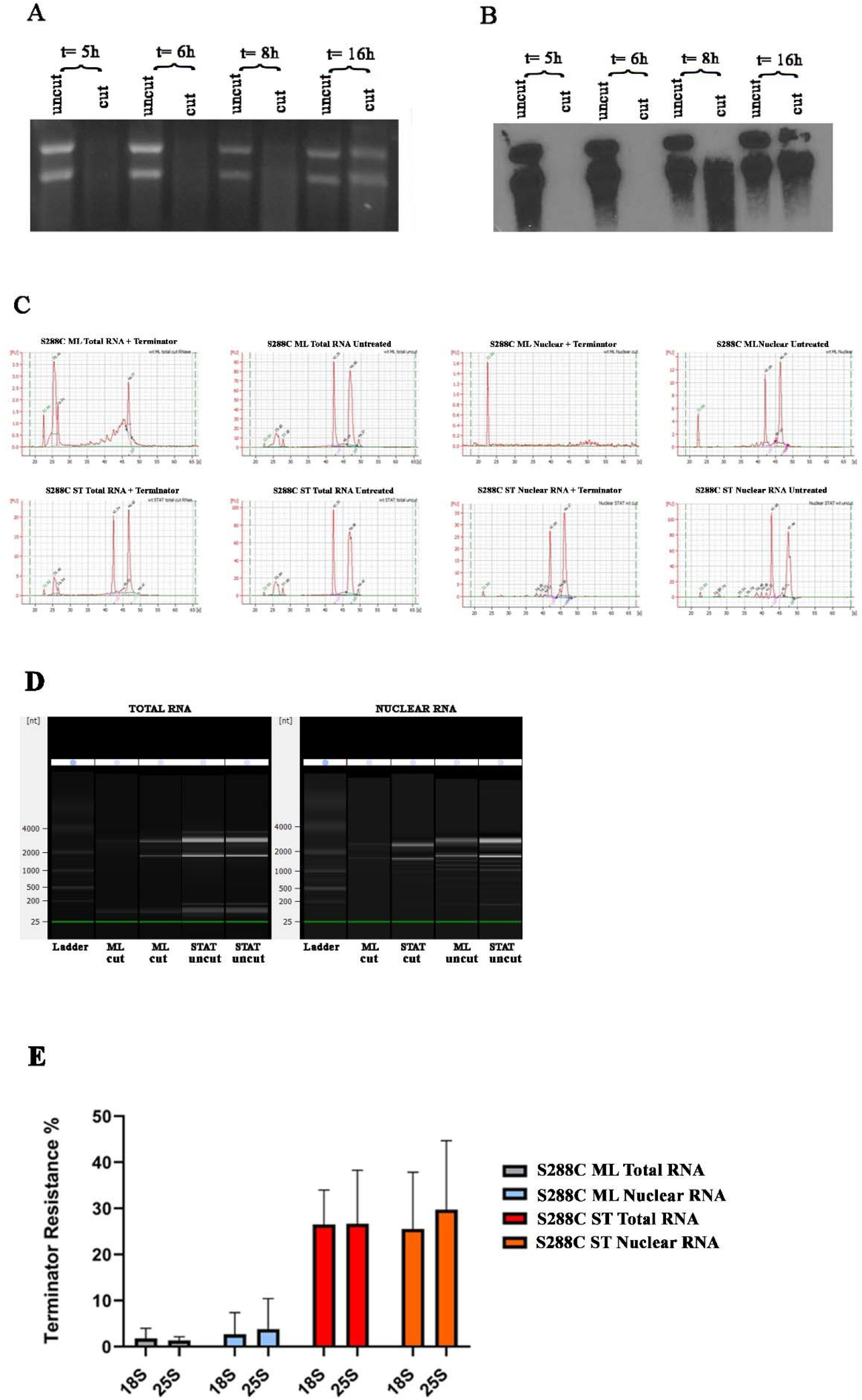
Terminator resistant 18S and 25S rRNA molecules in *Saccharomyces cerevisiae*. SYBR-gold stained gel (A) and Northern blot (B) showing rRNA extracted at different time points either treated (cut) or untreated (uncut) by Terminator. (C) Electropherograms used to validate the quality of RNA and to confirm the presence of Terminator resistant molecules. (D) Gel image generated from electropherograms by bioanalyzer software. (E) Terminator resistance percentage of ribosomal and nuclear RNA extracted from mid log (ML) and stationary (ST) wild type *S. cerevisiae*. Error bars represent standard deviation from three different experiments.

Figure 2 represents the Bioanalyzer assessment of ribosomal RNA from polymerase switched mutant *Saccharomyces cerevisiae.* As can be seen, both single and double mutant yeasts produce Terminator resistant 18S and 25S, (Fig 2A-B) with the double mutant producing them in the highest amount (Fig. C). This is true for both the total RNA and nuclear RNAs (Fig. 2C). In these mutants the percent of resistant 18S and 25S components from nuclear extracts are somewhat more than those in total RNA, again indicating that they are newly produced in the nucleolus and not modified in the cytoplasm. These data are important for several reasons. First, during the active growth phase in the wild type cells, where RNAP I is well established to be the transcribing polymerase, we do not detect these Terminator resistant molecules. This makes a role for RNAP I in their production unlikely. On the other hand, the fact that the double mutant, which has no functional RNAP I, only RNAP II for rRNA transcription, produces 18S and 25S at all and most efficiently, firmly establishes a role for RNAP II in the genesis of these molecules. The fact that the single mutant produces them in smaller amounts as compared to the double mutant, is also interesting. The single mutant has a functional RNAP I but its efficiency in gaining access to its promoter is limited due to the UAF component mutation. To whatever extent it can access its promoter, it will produce some Terminator sensitive 18S and 25S and reduce the percentage of Terminator resistant 18S and 25S produced by the cell. This is consistent with what we see in the single mutant.

**FIGURE 2.**
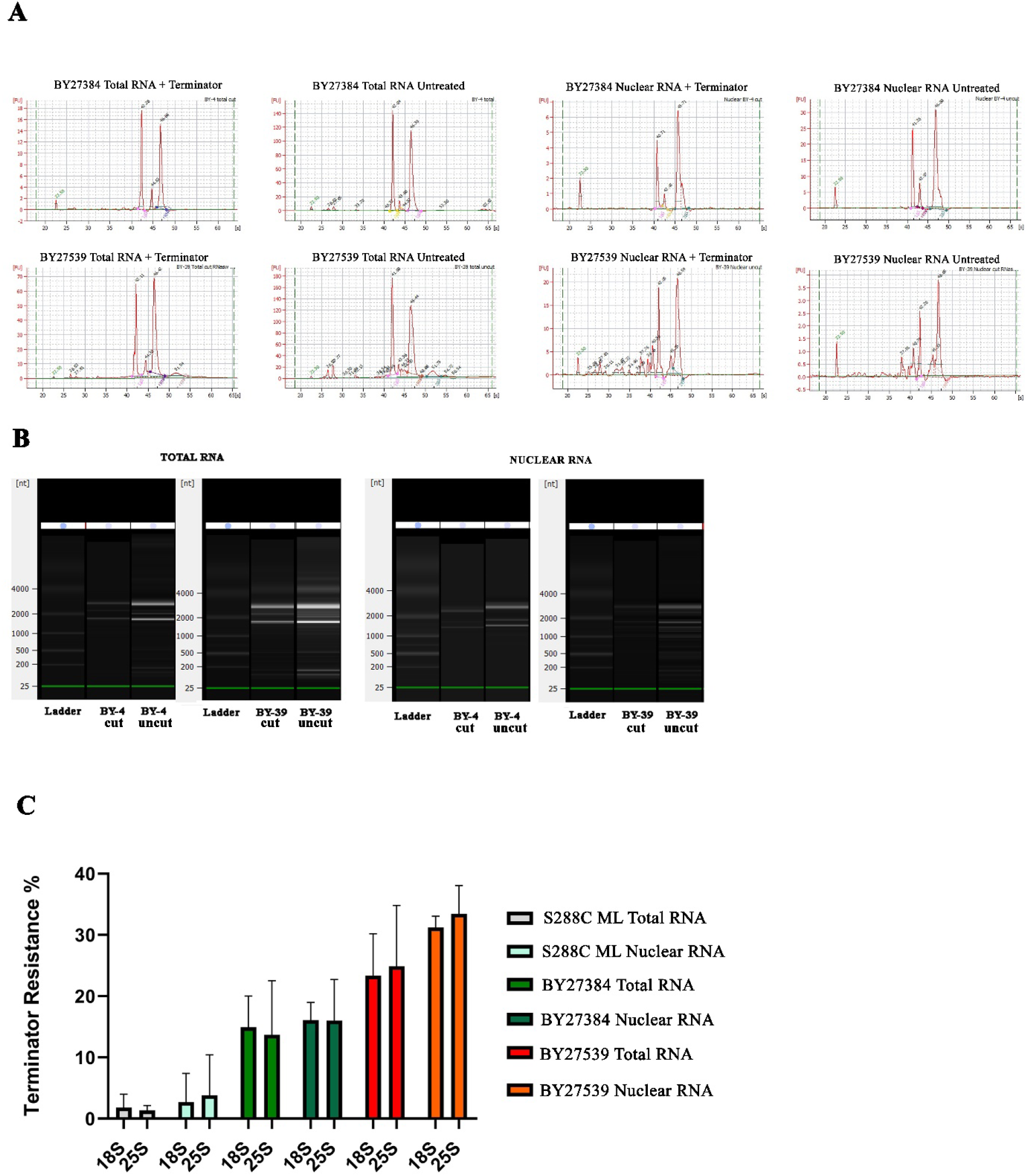
Terminator resistance of rRNA in single (BY27384) and double mutant (BY27539) *S. cerevisiae*. (A) Electropherograms used to validate the quality of RNA and to confirm the presence of Terminator resistant molecules. (B) Gel image generated from electropherograms by bioanalyzer software. (C) Terminator resistance percentage of ribosomal and nuclear RNA extracted from mid log (ML), single mutant (BY27384) and double mutant (BY27539) *S. cerevisiae*. Error bars represent standard deviation from three different experiments.

Resistance to Terminator digestion of 18S and 25S can arise in several ways. The single phosphate usually present at the 5’-end of these processed molecules can be removed or modified by addition to the single phosphate. This can include additional phosphate(s) with or without a cap structure or some other molecule. Data shown in Figure 3 indicates that the addition of one or more phosphate is the basis of the resistance. Terminator resistant RNAs isolated from yeast in stationary phase or from the mutants are made susceptible to Terminator by first digesting them with a decapping enzyme. These enzymes remove the cap structures from RNA by cutting between phosphates and can leave a single phosphate on the 5’-end of the RNAs making them susceptible to elimination by Terminator. This indicates, that at the minimum these Terminator resistant 18S and 25S molecules have more than one phosphate at their 5’-end.

**FIGURE 3.**
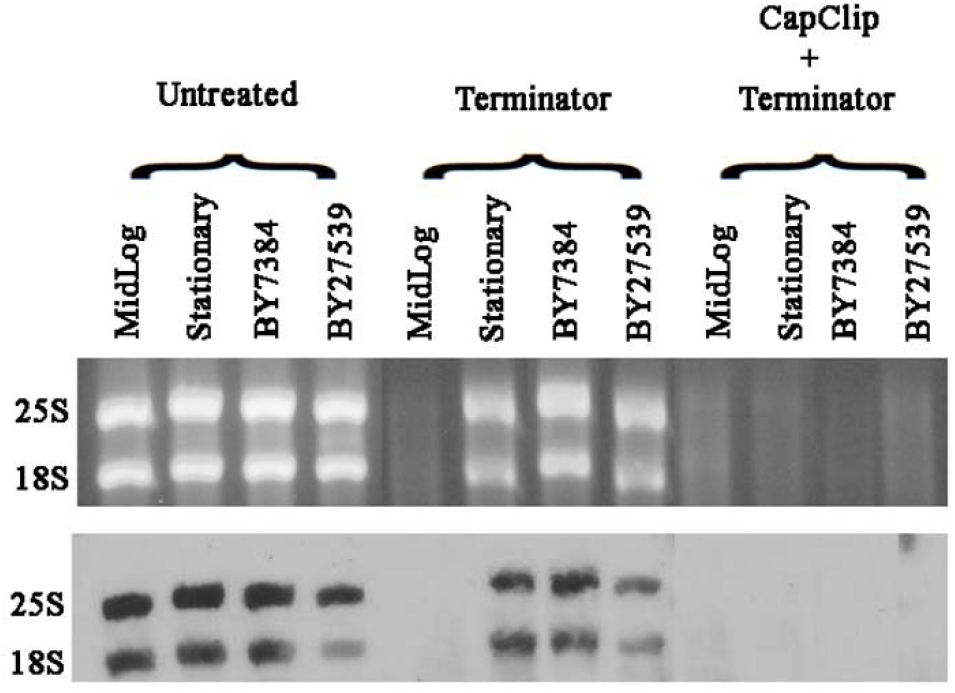
5’end analysis of 18S and 25S molecules in wild type and mutant *S. cerevisiae*. SYBR-gold stained gel and Northern blot shows the effect decapping followed by Terminator treatment on rRNA molecules 18S and 25S.

**FIGURE 4.**
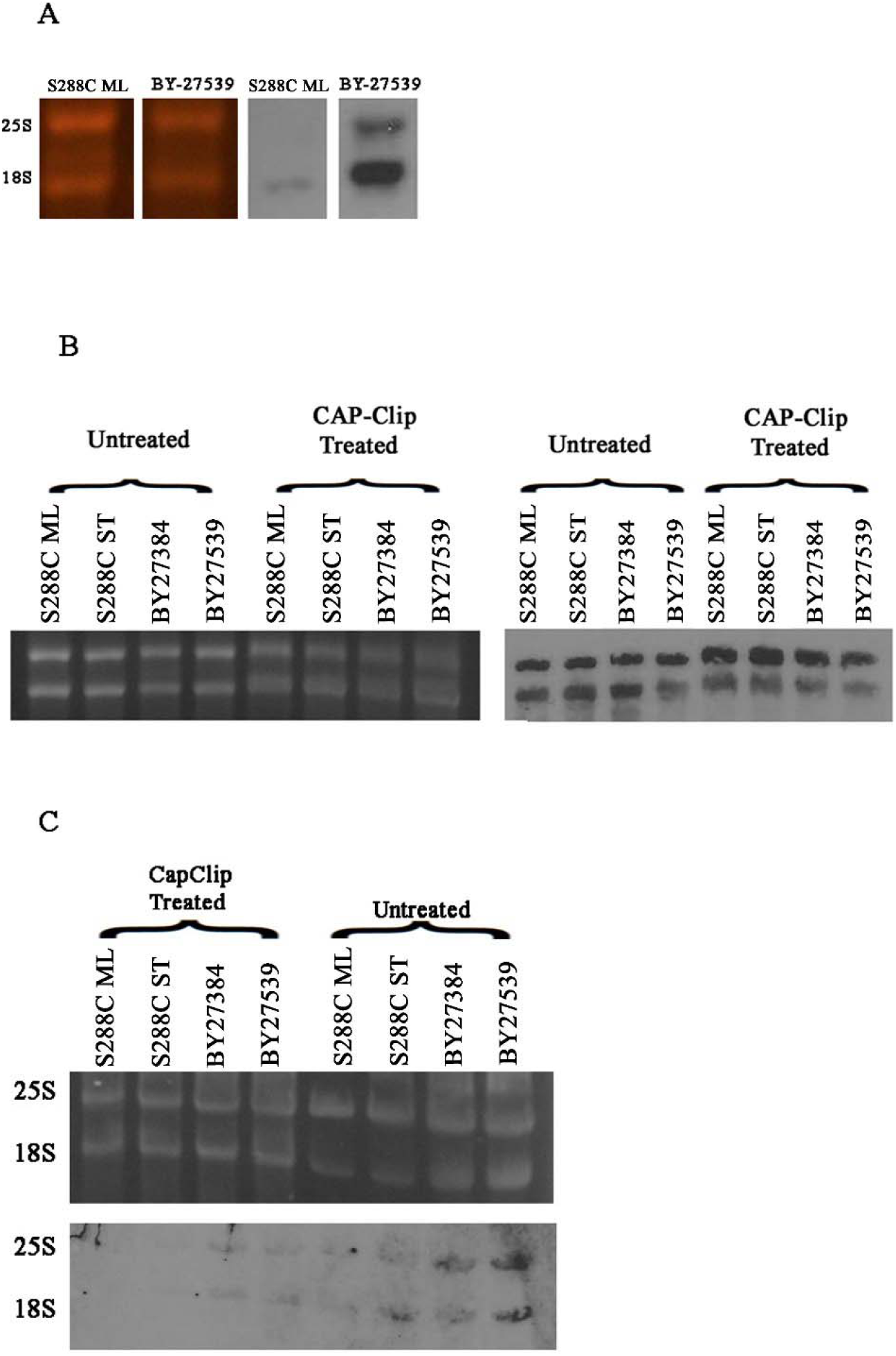
5’-cap analysis. (A) SYBR-gold stained gel and immunoblot using cap-specific antibody H20 indicating presence of cap in mutant and stationary wild type S. cerevisiae. (B) SYBR-gold stained gel and Northern blot with 18S and 25S probes showing rRNA that has been treated with CAP-Clip or untreated. Decapping enzyme does not degrade RNA. (C) Gel and corresponding Immunoblot using H20 antibody. Decapping clearly decreased signal generated by the antibody, with the greatest difference in the double mutant.

To see if any cap structure is present on these molecules we utilized the trimethyl cap monoclonal antibody H20, widely used in cap detection studies (Bochnig et al., 1987). As can be seen in Figure 3A the antibody strongly reacts with 18S and 25S rRNAs derived from the double mutant and not at all or only minimally from molecules isolated from wild type yeast in active growth phase. The minimal signal from wild type yeast RNA likely represents non-specific binding by the antibody. The stained gels show that the difference in intensity on the immunoblot is not related to differences in amount of RNA present. Decapping these molecules (Fig. 3C) decreases their immunoblot signals, again confirming the presence of a cap on the phosphates. A Northern blot (Fig. 3B) shows that the decapping enzyme does not degrade the RNA and therefore is not the reason for the decrease in immunoblot intensity.

These data indicate that RNAP II is producing 18S and 25S RNA molecules with capped tri-phosphorylated 5’-ends. Such molecules would be predictably resistant to Terminator digestion. Interestingly, polyadenylation of mRNAs is a has also been reported for rRNA (Fleischmann and Liu, 2001); (Fleischmann et al., 2004); (Kuai et al., 2004); (Jenjaroenpun et al., 2018) which fits well with our data. The mechanism by which this polymerase produces these molecules is unknown and will likely to be a novel addition to RNAP II’s activities. The fact that a newly transcribed RNAP II product gets modified by capping is not unusual, as it happens to all known RNA II transcripts, including mRNAs, miRNAs, lncRNAs, snoRNAs and snRNAs (Rambout and Maquat, 2020). It is also well established that RNAP II can be involved in rRNA production in a polycistronic fashion. It has been shown to gain access to rDNA promoter site with nutritional deprivation (Vallabhaneni, 2015). As mentioned in the introduction this has also been shown to be true in the polymerase switched mutants. These polycistronic transcripts are processed into 18S and 25S components with a single phosphate at the 5’-end and therefore susceptible to be eliminated by Terminator. The fact that Terminator eliminates over 50% of 18S and 25S confirms this. What is unknown is, how the rest of these RNA molecules with cap-associated Terminator resistance are produced.

Two modalities can be postulated each with its challenges. As all polymerases initiate transcription with nucleoside triphosphates (NTPs), in general an RNA molecule with three phosphates at its 5’-end, can be newly transcribed and if capped by, RNAP II. In *Candida albicans*, with oligo-ligation and primer extension, we have shown that the extra phosphates on these resistant molecules were at the processing site both for 18S and 25S (Fleischmann and Rocha, 2018). While we have not repeated this with *Saccharomyces cerevisiae*, being that their sizes are that of the processed molecules, makes it likely for this yeast too. Thus, if these molecules are newly transcribed, then in addition to its polycistronic transcription, RNAP II would also be initiating 18S and 25S specific transcriptions. There are no canonical sequences for an RNAP II associated promoter immediately upstream of 18S and 25S. It would have to be an unusual promoter capable of attracting all the transcription factors required by RNAP II for initiation of transcription.

The other modality would involve the capping of processed 18S and 25S molecules initially transcribed in a polycistronic manner. Capping of pre-mRNAs in *Saccharomyces cerevisiae*, is carried out co-transcriptionally by the capping enzyme complex (CE) made up of the triphosphatase Cet1 and the guanylyltransferase Ceg1 (Bharati et al., 2016). It is well established in yeast, that the capping enzyme complex interacts with the polymerase subunit of RNAP II at the C-terminal heptad repeats (Gu et al., 2010). In fact, even the third enzyme involved with capping, namely N7 methyltransferase also interacts with the C-terminal repeats (Ramanathan et al., 2016). As processing of polycistronic rRNA can occur co-transcriptionally, if RNAP II is the transcribing polymerase of these 18S and 25S molecules, the capping machinery would also be available co-transcriptionally. The difficulty with this scenario is that the Cet1 triphosphatase requires three phosphates as substrates at the 5’-position. Thus, there would have to be a kinase present in the nucleus capable in adding co-transcriptionally two phosphates to the single 5’-phosphate of the processed 18S and 25S molecules. There are examples that point to such a possibility. Cytoplasmic recapping of mRNAs has been shown in mammalian cells. Nudix family decapping enzymes such as DCP2, cleave between the α and β phosphates of the tri-phosphate cap leaving a 5’-monophosphate RNA in the cytoplasm. The nuclear mammalian capping complex RNGTT, is present in the cytoplasm also and for its triphosphates to function in recapping the 5’-monophosphate decapped mRNAs, they would need to have phosphates added to their 5’-end. Cytoplasmic capping enzyme complex with such kinase activity has been shown to be present in mammalian cells (Otsuka et al., 2009) (Trotman and Schoenberg, 2019). Similarly, in the kinetoplastid *Trypanosoma brucei*, a cytoplasmic guanylyltransferase with 5’-RNA kinase activity capable of transforming a pRNA to a ppRNA, allowing a GMP transfer from GTP has been described (Ignatochkina et al., 2015)

In *Candida albicans* we have been able to detect Terminator resistant 18S and 25S RNAs in ribosomes isolated from stationary yeast indicating that they are functional (Fleischmann and Rocha, 2018). A potential for such degradation resistant molecules for the cell would be to maintain the protein producing capacity of the cell under nutritional duress. Our data indicates that this new role for RNAP II is not limited to mutational limitations of RNAP I but is present in wild type organisms during some part of the growth cycle, giving RNAP II an additional role in ribosomal production.

## References

Abraham, K.J., Khosraviani, N., Chan, J.N.Y., Gorthi, A., Samman, A., Zhao, D.Y., Wang, M., Bokros, M., Vidya, E., Ostrowski, L.A., et al. (2020). Nucleolar RNA polymerase II drives ribosome biogenesis. Nature 585, 298–302.

Bharati, A.P., Singh, N., Kumar, V., Kashif, M., Singh, A.K., Singh, P., Singh, S.K., Siddiqi, M.I., Tripathi, T., and Akhtar, M.S. (2016). The mRNA capping enzyme of Saccharomyces cerevisiae has dual specificity to interact with CTD of RNA Polymerase II. Sci Rep 6, 31294.

Bochnig, P., Reuter, R., Bringmann, P., and Luhrmann, R. (1987). A monoclonal antibody against 2,2,7-trimethylguanosine that reacts with intact, class U, small nuclear ribonucleoproteins as well as with 7-methylguanosine-capped RNAs. Eur J Biochem 168, 461–467.

Clark-Walker, G.D., and Azad, A.A. (1980). Hybridizable sequences between cytoplasmic ribosomal RNAs and 3 micron circular DNAs of Saccharomyces cerevisiae and Torulopsis glabrata. Nucleic Acids Res 8, 1009–1022.

Conrad-Webb, H., and Butow, R.A. (1995). A polymerase switch in the synthesis of rRNA in Saccharomyces cerevisiae. Mol Cell Biol 15, 2420–2428.

Fleischmann, J., and Liu, H. (2001). Polyadenylation of ribosomal RNA by Candida albicans. Gene 265, 71–76.

Fleischmann, J., Liu, H., and Wu, C.P. (2004). Polyadenylation of ribosomal RNA by Candida albicans also involves the small subunit. BMC Mol Biol 5, 17.

Fleischmann, J., and Rocha, M.A. (2018). Nutrient depletion and TOR inhibition induce 18S and 25S ribosomal RNAs resistant to a 5’-phosphate-dependent exonuclease in Candida albicans and other yeasts. BMC Mol Biol 19, 1.

Fleischmann, J., Rocha, M.A., Hauser, P.V., Gowda, B.S., and Pilapil, M.G.D. (2020). Exonuclease resistant 18S and 25S ribosomal RNA components in yeast are possibly newly transcribed by RNA polymerase II. BMC Mol Cell Biol 21, 59.

Goetze, H., Wittner, M., Hamperl, S., Hondele, M., Merz, K., Stoeckl, U., and Griesenbeck, J. (2010). Alternative chromatin structures of the 35S rRNA genes in Saccharomyces cerevisiae provide a molecular basis for the selective recruitment of RNA polymerases I and II. Mol Cell Biol 30, 2028–2045.

Gu, M., Rajashankar, K.R., and Lima, C.D. (2010). Structure of the Saccharomyces cerevisiae Cet1-Ceg1 mRNA capping apparatus. Structure 18, 216–227.

Haag, J.R., and Pikaard, C.S. (2007). RNA polymerase I: a multifunctional molecular machine. Cell 131, 1224–1225.

Ignatochkina, A.V., Takagi, Y., Liu, Y., Nagata, K., and Ho, C.K. (2015). The messenger RNA decapping and recapping pathway in Trypanosoma. Proc Natl Acad Sci U S A 112, 6967–6972.

Jager, D., Sharma, C.M., Thomsen, J., Ehlers, C., Vogel, J., and Schmitz, R.A. (2009). Deep sequencing analysis of the Methanosarcina mazei Go1 transcriptome in response to nitrogen availability. Proc Natl Acad Sci U S A 106, 21878–21882.

Jenjaroenpun, P., Wongsurawat, T., Pereira, R., Patumcharoenpol, P., Ussery, D.W., Nielsen, J., and Nookaew, I. (2018). Complete genomic and transcriptional landscape analysis using third-generation sequencing: a case study of Saccharomyces cerevisiae CEN.PK113-7D. Nucleic Acids Res 46, e38.

Kargas, V., Castro-Hartmann, P., Escudero-Urquijo, N., Dent, K., Hilcenko, C., Sailer, C., Zisser, G., Marques-Carvalho, M.J., Pellegrino, S., Wawiorka, L., et al. (2019). Mechanism of completion of peptidyltransferase centre assembly in eukaryotes. Elife 8.

Keener, J., Dodd, J.A., Lalo, D., and Nomura, M. (1997). Histones H3 and H4 are components of upstream activation factor required for the high-level transcription of yeast rDNA by RNA polymerase I. Proc Natl Acad Sci U S A 94, 13458–13462.

Kuai, L., Fang, F., Butler, J.S., and Sherman, F. (2004). Polyadenylation of rRNA in Saccharomyces cerevisiae. Proc Natl Acad Sci U S A 101, 8581–8586.

Nogi, Y., Vu, L., and Nomura, M. (1991). An approach for isolation of mutants defective in 35S ribosomal RNA synthesis in Saccharomyces cerevisiae. Proc Natl Acad Sci U S A 88, 7026–7030.

Oakes, M., Siddiqi, I., Vu, L., Aris, J., and Nomura, M. (1999). Transcription factor UAF, expansion and contraction of ribosomal DNA (rDNA) repeats, and RNA polymerase switch in transcription of yeast rDNA. Mol Cell Biol 19, 8559–8569.

Otsuka, Y., Kedersha, N.L., and Schoenberg, D.R. (2009). Identification of a cytoplasmic complex that adds a cap onto 5’-monophosphate RNA. Mol Cell Biol 29, 2155–2167.

Planta, R.J., and Raue, H.A. (1988). Control of ribosome biogenesis in yeast. Trends Genet 4, 64–68.

Ramanathan, A., Robb, G.B., and Chan, S.H. (2016). mRNA capping: biological functions and applications. Nucleic Acids Res 44, 7511–7526.

Rambout, X., and Maquat, L.E. (2020). The nuclear cap-binding complex as choreographer of gene transcription and pre-mRNA processing. Genes Dev 34, 1113–1127.

Schroeder, A., Mueller, O., Stocker, S., Salowsky, R., Leiber, M., Gassmann, M., Lightfoot, S., Menzel, W., Granzow, M., and Ragg, T. (2006). The RIN: an RNA integrity number for assigning integrity values to RNA measurements. BMC Mol Biol 7, 3.

Siddiqi, I.N., Dodd, J.A., Vu, L., Eliason, K., Oakes, M.L., Keener, J., Moore, R., Young, M.K., and Nomura, M. (2001). Transcription of chromosomal rRNA genes by both RNA polymerase I and II in yeast uaf30 mutants lacking the 30 kDa subunit of transcription factor UAF. EMBO J 20, 4512–4521.

Thiry, M., and Lafontaine, D.L. (2005). Birth of a nucleolus: the evolution of nucleolar compartments. Trends Cell Biol 15, 194–199.

Trotman, J.B., and Schoenberg, D.R. (2019). A recap of RNA recapping. Wiley Interdiscip Rev RNA 10, e1504.

Turowski, T.W., and Tollervey, D. (2016). Transcription by RNA polymerase III: insights into mechanism and regulation. Biochem Soc Trans 44, 1367–1375.

Vallabhaneni, A.R. (2015). Saccharomyces cerevisiae synthesizes ribosomal RNA using RNA polymerase II during nitrogen deprivation (Texas Woman’s University).

Vu, L., Siddiqi, I., Lee, B.S., Josaitis, C.A., and Nomura, M. (1999). RNA polymerase switch in transcription of yeast rDNA: role of transcription factor UAF (upstream activation factor) in silencing rDNA transcription by RNA polymerase II. Proc Natl Acad Sci U S A 96, 4390–4395.

Warner, J.R. (1999). The economics of ribosome biosynthesis in yeast. Trends Biochem Sci 24, 437–440.

Woolford, J.L., Jr., and Baserga, S.J. (2013). Ribosome biogenesis in the yeast Saccharomyces cerevisiae. Genetics 195, 643–681.

Yano, R., and Nomura, M. (1991). Suppressor analysis of temperature-sensitive mutations of the largest subunit of RNA polymerase I in Saccharomyces cerevisiae: a suppressor gene encodes the second-largest subunit of RNA polymerase I. Mol Cell Biol 11, 754–764.

